# Towards Targeting the Disordered SARS-CoV-2 Nsp2 C-terminal Region: Partial Structure and Dampened Mobility Revealed by NMR Spectroscopy

**DOI:** 10.1101/2020.11.09.374173

**Authors:** Miguel Mompeán, Miguel Á. Treviño, Douglas V. Laurents

## Abstract

Intrinsically disordered proteins (IDPs) play essential roles in regulating physiological processes in eukaryotic cells. Many virus use their own IDPs to “hack” these processes to disactive host defenses and promote viral growth. Thus, viral IDPs are attractive drug targets. While IDPs are hard to study by X-ray crystallography or cryo-EM, atomic level information on their conformational perferences and dynamics can be obtained using NMR spectroscopy. SARS-CoV-2 Nsp2 interacts with human proteins that regulate translation initiation and endosome vesicle sorting, and the C-terminal region of this protein is predicted to be disordered. Molecules that block these interactions could be valuable leads for drug development. To enable inhibitor screening and to uncover conformational preferences and dynamics, we have expressed and purified the ^13^C,^15^N-labeled C-terminal region of Nsp2. The ^13^Cβ and backbone ^13^CO, ^1^HN, ^13^Cα and ^15^N nuclei were assigned by analysis of a series of 2D ^1^H-^15^N HSQC and ^13^C-^15^N CON as well as 3D HNCO, HNCA, CBCAcoNH and HncocaNH spectra. Overall, the chemical shift data confirm that this region is chiefly disordered, but contains two five-residue segments that adopt a small population of β-strand structure. Whereas the region is flexible on ms/ms timescales as gauged by T_1ρ_ measurements, the {^1^H}-^15^N NOEs reveal a flexibility on ns/ps timescales that is midway between a fully flexible and a completely rigid chain.

## Introduction

The pandemic provoke by Sars-CoV-2 is sparking unprecedented efforts in the development of vaccine. Nevertheless, drugs which block the activity of viral proteins should also be sought, since Sars-CoV-2 vaccines may well lack availability, public acceptance and complete efficacy, particularly for the immune compromised. Intrinsically disordered proteins (IDPs) play essential roles in regulating physiological processes in eukaryotic cells by forming weak protein/protein interactions. Many virus use their own IDPs to “hack” these processes to disactive host defenses and promote viral growth (Davey *et al.*, 2011). Thus, viral IDPs are attractive drug targets. While the flexibility of IDPs generally precludes their study by X-ray crystallography or cryo-EM, atomic level information on their conformational preferences and dynamics can be obtained using NMR spectroscopy.

Using bioinformatics tools, a C-terminal region of SARS-CoV-2 Nsp2 has been predicted to be disordered (Giri *et al.*, 2020). Interestingly, this Nsp2 region appears to be more disordered in SARS-CoV-2 than in its close homologs human SARS (responsible for the 2003 outbreak) and bat coronavirus (see Sup. Fig. 9 in Giri *et al.*, 2020). A recent study (Gordon *et al.*, 2020) of the SARS-CoV-2 interactome reported that Nsp2 interacts with human proteins GIGYF2 and EIF4E2, which regulate translation initiation, as well as WASHC4 and FKBP15, which are involved in endosome vesicle sorting. Nsp2 from human SARS was also reported to interact with Prohibitin1/2, a cell proliferation modulator with tumor suppression activity (Cornilley-Ty *et al.*, 2009). Molecules that block these interactions could be valuable leads for drug development. The main objective of this study is to characterize the conformational preferences and dynamics of the putatively disordered C-terminal region of SARS CoV-2 Nsp2 (Nsp2 CtDR) using NMR spectroscopy. The assignments obtained could also guide future studies of interactions between Nsp2, human target proteins and inhibitors.

## Materials and Methods

### Bioinformatics

The algorithm PrDOS (Ishida & Kinoshita, 2007) was used to predict the disordered regions in the Nsp2 sequence. The default setting for 5% false positives was used. This method consists of two predictors, one based on the local sequence composition and a second based on the level of sequence conservation in homologous proteins.

### Sample production and isotopic labeling

To enable inhibitor screening and to uncover conformational preferences and dynamics, we have expressed and purified the ^13^C,^15^N-labeled C-terminal region of Nsp2. The full production methodology will be reported soon [Covid19-NMR Consortium, 2020]. Briefly, the segment K_55_7EIIFLEGETLPTEVLTEEVVLKTGDLQPLEQPTSEAVEAPLVGT_601_, from the C-terminus of Nsp2 was expressed in minimal media containing ^15^NH_4_Cl and ^13^C-glucose as the sole sources of nitrogen and carbon, respectively, and the domain was purified using the His-tag / Ni^++^ column chromatography approach.

### NMR spectroscopy

Samples for NMR spectrscopy contained ^13^C,^15^N-labeled Nsp2 CtDR at pH 6.3 in 5.0 mM Na_2_HPO_4_ / NaH_2_PO_4_ buffer with 10.0 mM NaCl, 10% D_2_O and sodium trimethylsilypropanesulfonate (DSS) as the internal chemical shift reference. Two series of spectra were recorded at 5 °C, one with and one without the presence of an N-terminal H-tag whose sequence is MAHHHHHHGTGTGSNDDDD-K.

A series of 2D ^1^H-^15^N HSQC and ^13^C-^15^N CON as well as 3D HNCO, HNCA, CBCAcoNH and HncocaNH (Pantoja-Uceda & Santoro, 2009) spectra were recorded on a Bruker Neo 800 MHz (^1^H) spectrometer equipped with a cryoprobe and z-gradients. The NMR spectral parameters are listed in **Table 1**. The spectra were transformed with Topspin 4.0.8 (Bruker Biospin) and were assigned manually with the aid of the program Sparky (Lee *et al.*, 2015). In addition, a series of 2D ^1^H-^15^N HSQC-based experiments were recorded to determine the {^1^H}-^15^N NOE and R_1ρ_ relaxation rates in order to assess dynamics on ps/ns and μs/ms timescales, respectively. These spectra were recorded without non-uniform sampling and were transformed without applying linear prediction. Peak integrals were measured using Topspin 4.0.8. R_1ρ_ relaxation rates were obtained by using the program Kaleidagraph (Synergy Software, version 3.6) to fit an exponential equation: I(t) = Io· exp(−k·t) + I∞, where I(t) is the peak integral at time t, k is the rate, and I∞ is the intensity at infinite time. The R_1ρ_ uncertainties reported here are those obtained from the least squares fit. In the case of the {^1^H}-^15^N NOE ratios, the uncertainties were calculated as the ratio of the noise (estimated as the standard deviation of the integral of several areas lacking peaks) times √2 to the peak integral measured without applying the NOE.

**Table 1:** NMR Spectral Parameters

| **Experiment** | **Number of Scans** | **Sweep Width (ppm)** | **Matrix** |
| --- | --- | --- | --- |
| **Assignment** |  |  |  |
| 1D $^1\text{H}$ | 48 | 12 | 32k |
| 2D $^1\text{H}$-$^{15}\text{N}$ HSQC | 4 | 11 $^1\text{H}$ x 20 $^{15}\text{N}$ | 2k x 256 |
| 2D CON | 8 | 11 $^{13}\text{CO}$ x 34 $^{15}\text{N}$ | 1k x 512 |
| 3D HNCO | 8 | 11 $^1\text{H}$ x 20 $^{15}\text{N}$ x 12 $^{13}\text{C}$ | 2k x 48 x 256 |
| 3D HNCA | 8 | 11 $^1\text{H}$ x 20 $^{15}\text{N}$ x 25 $^{13}\text{C}_\alpha$ | 2K x 48 x 128 |
| 3D CBCAcoNH | 8 | 2k $^1\text{H}$ x 20 $^{15}\text{N}$ x 55 $^{13}\text{C}$ | 2k x 48 x 128 |
| 3D HncocaNH | 8 | 10 $^1\text{H}$ x 20 $^{15}\text{N}$ x 10 $^1\text{H}$ | 2k x 64 x 64 |
| **Relaxation** |  |  |  |
| 2D HSQCTEFT3GPSI<br>($T_{1\rho}$)<sup>*</sup> | 8 | 10 $^1\text{H}$ x 20 $^{15}\text{N}$ | 2k x 384 |
| 2D HSQCNOEFGPSI<br>{$^1\text{H}$}-$^{15}\text{N}$ NOE<sup>‡</sup> | 16 | 10 $^1\text{H}$ x 20 $^{15}\text{N}$ | 2k x 384 |

<sup>\*</sup> Delays used (in seconds): 0.008, 0.300, 0.036, 0.076, 0.900, 0.150, 0.500, 1.350.
<sup>‡</sup> Recycle delay = 10 seconds.

## Results

### 1. Bioinformatics analysis

The C-terminal region of Nsp2 was recently predicted to be disordered by Giri *et al.*, 2020, who applied the PONDR^®^ and IUPRED algorithms. Nevertheless, this region contains a rather high proportion of hydrophobic residues which generally promote order and folding. To corroborate these findings, we have applied the same tools as Giri *et al.*, 2020 as well as the PrDOS disorder prediction algorithm (Ishida & Kinoshita, 2007) to the Nsp2 sequence (**Figure 1**). Whereas the results show that the C-terminal region spanning approximately residues 555 – 600 have a high disorder propensity as compared to the rest of the protein, the score falls short of the threshold for some algorithms used by Giri *et al.*, (2020) (Fig. 1B) and is close to the threshold for PrDOSfor predicting disordered (**Figure 1**). Considering this finding, and the fact that disorder prediction programs are sometimes wrong (Treviño *et al*, 2018), we have used NMR spectroscopy to characterize structurally the putatively disordered region of Nsp2.

**Figure 1A.**
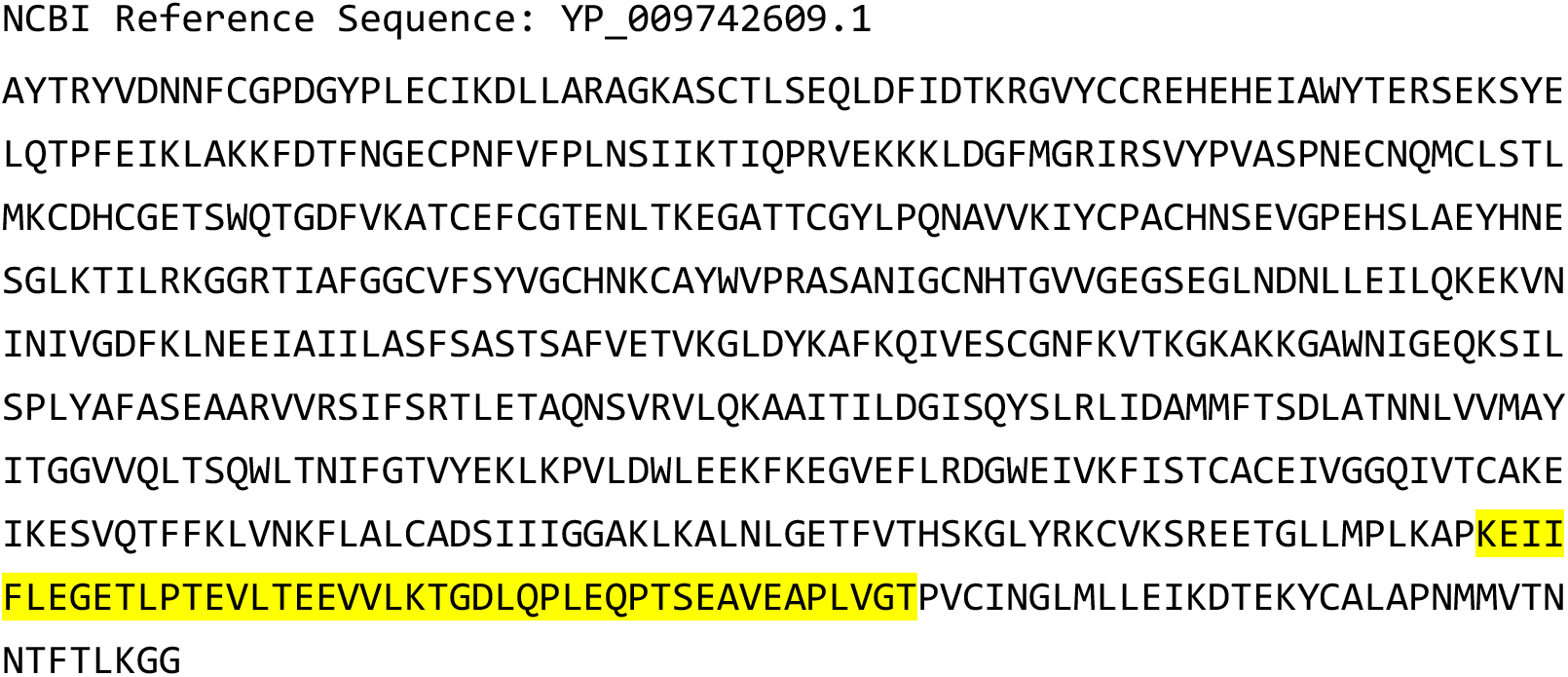
SARS-CoV-2 Nsp2 Primary Structure and Predicted Disordered Region. nsp2 [Severe acute respiratory syndrome coronavirus 2] The region examined here is highlighted in yellow.

**Figure 1B:**
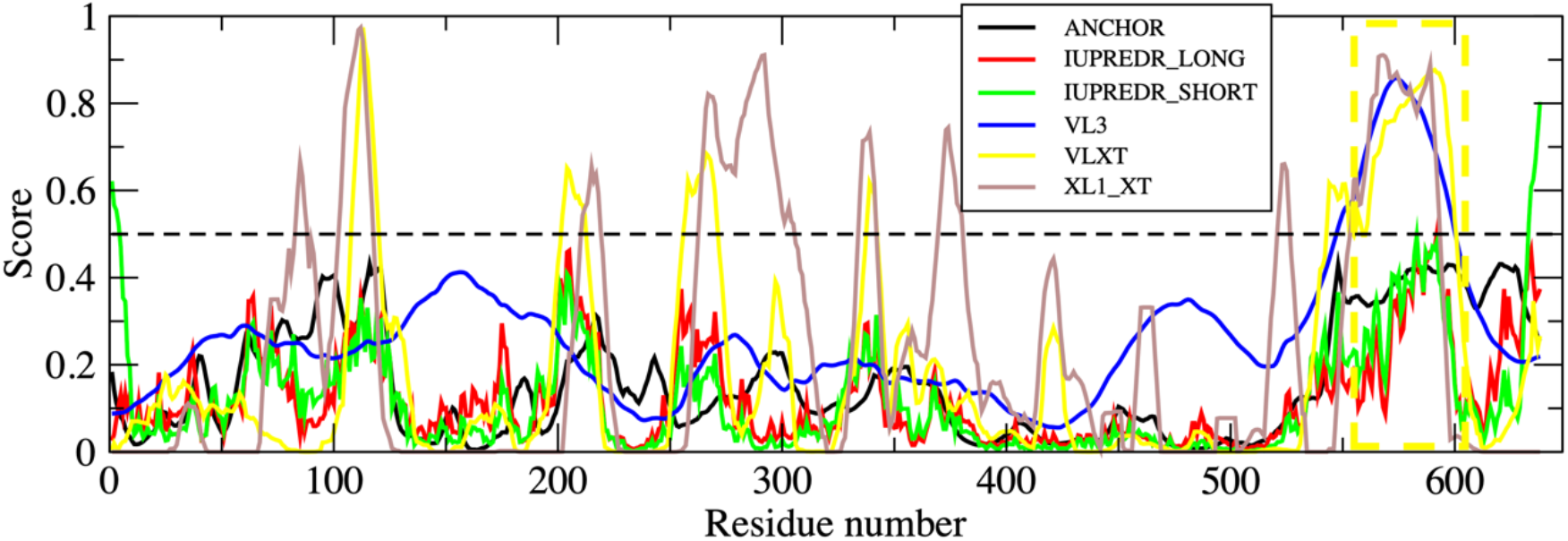
SARS-CoV-2 Nsp2 Primary Structure and Predicted Disordered Region. The disorder prediction obtained with a group of predictors used by Giri *et al.*, (2020) The dotted yellow box marks the region examined here.

**1C.**
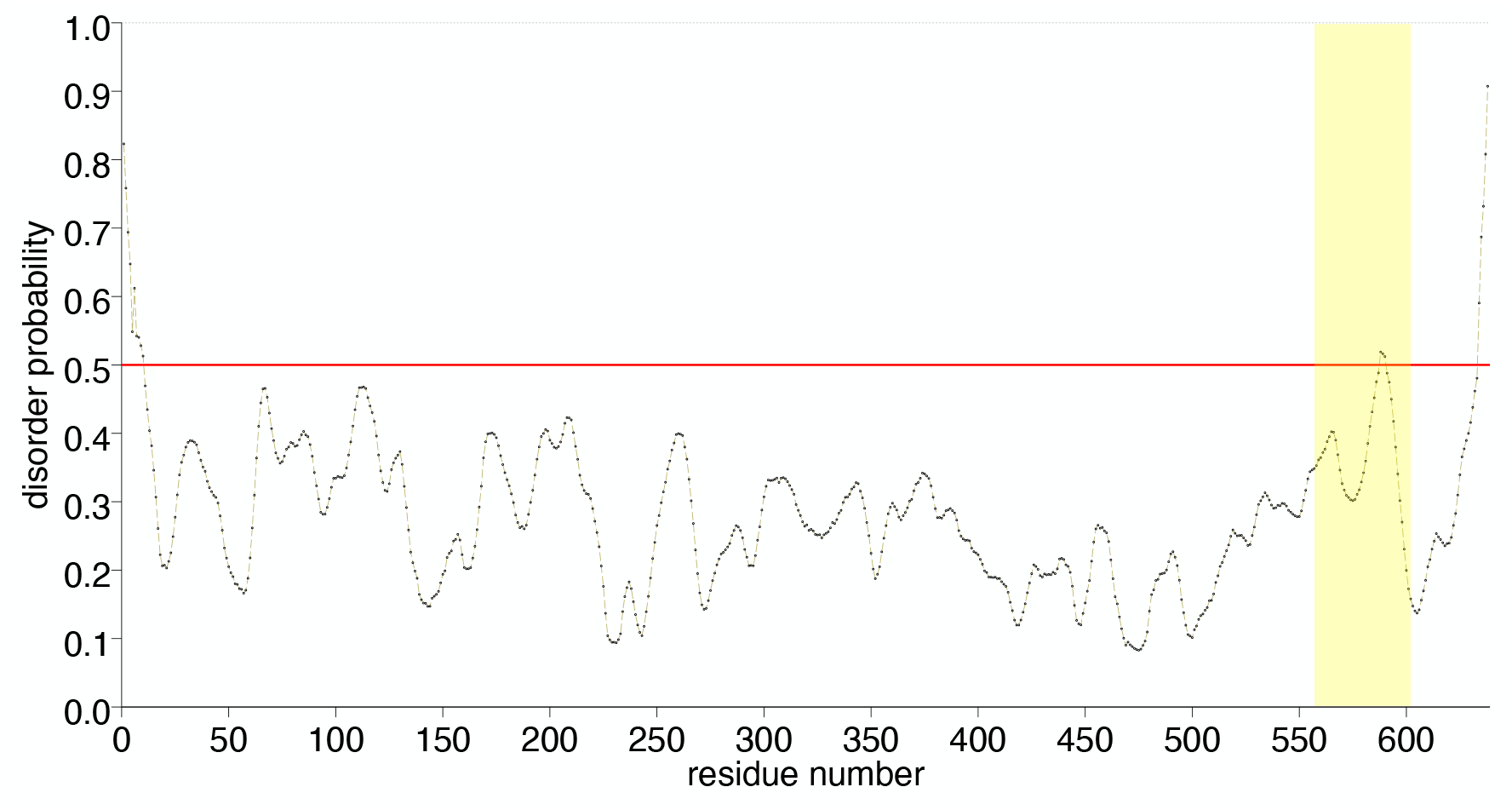
SARS-CoV-2 Nsp2 Primary Structure and Predicted Disordered Region. The disorder prediction obtained with PrDOS (Ishida & Kinoshita, 2007). The region of interest is shaded yellow.

### 2. NMR assignment and assessment of partial structure

The ^13^Cβ and backbone ^13^CO, ^1^HN, ^13^Cα and ^15^N nuclei were assigned by analysis of a series of 2D ^1^H-^15^N HSQC and ^13^C-^15^N CON as well as 3D HNCO, HNCA, CBCAcoNH and HncocaNH (Pantoja-Uceda & Santoro, 2009) spectra. The assignments are 98% ^13^Cα, 87% ^13^Cβ, 84% ^13^CO, 98% ^15^N and 98% ^1^HN complete for the region without the His tag; and 82% ^13^Cα, 82% ^13^Cβ, 82% ^13^CO, 91% ^15^N and 100% ^1^HN complete for the region without the His tag.

The assigned 2D ^1^H-^15^N HSQC spectra of Nsp2 CtDR with and without an N-terminal His tag are shown in **Figure 2**. The low ^1^HN signal dispersion suggests that the Nsp2 CtDR is disordered. The ^13^Cα, ^13^Cβ and ^13^CO conformational chemical shifts (Δδ) values are shown in **Figure 3**. Overall, the Δδ values are low, which evinces that the Nsp2 CtDR is disordered. However, two five-residues stretches spanning residues E_570_-VLTE_574_ and S_591_EAVE_595_ tend to adopt β-stands. The populations of these β-strands within the conformational ensemble is about 10%. In addition, a few discontinuous residues at the N-terminus of this region also show some tendency to adopt β-stand conformations (**Figure 3**). It is important to point out that the sample whose His tag was cleaved contained an additional proteolytic cleavage between residues K_579_-T_580_, which was identified on the basis of chemical shift alterations, the absence of an ^1^H-^15^N crosspeak for T_580_ and the relaxation measurements reported below. This break occurs between the two β-strands mentioned above. Since the β-strand populations for E_570_-E_574_ and S_591_-E_595_ are similar in both samples, this suggests that no interaction between these β-strands is necessary for them to form.

**Figure 2:**
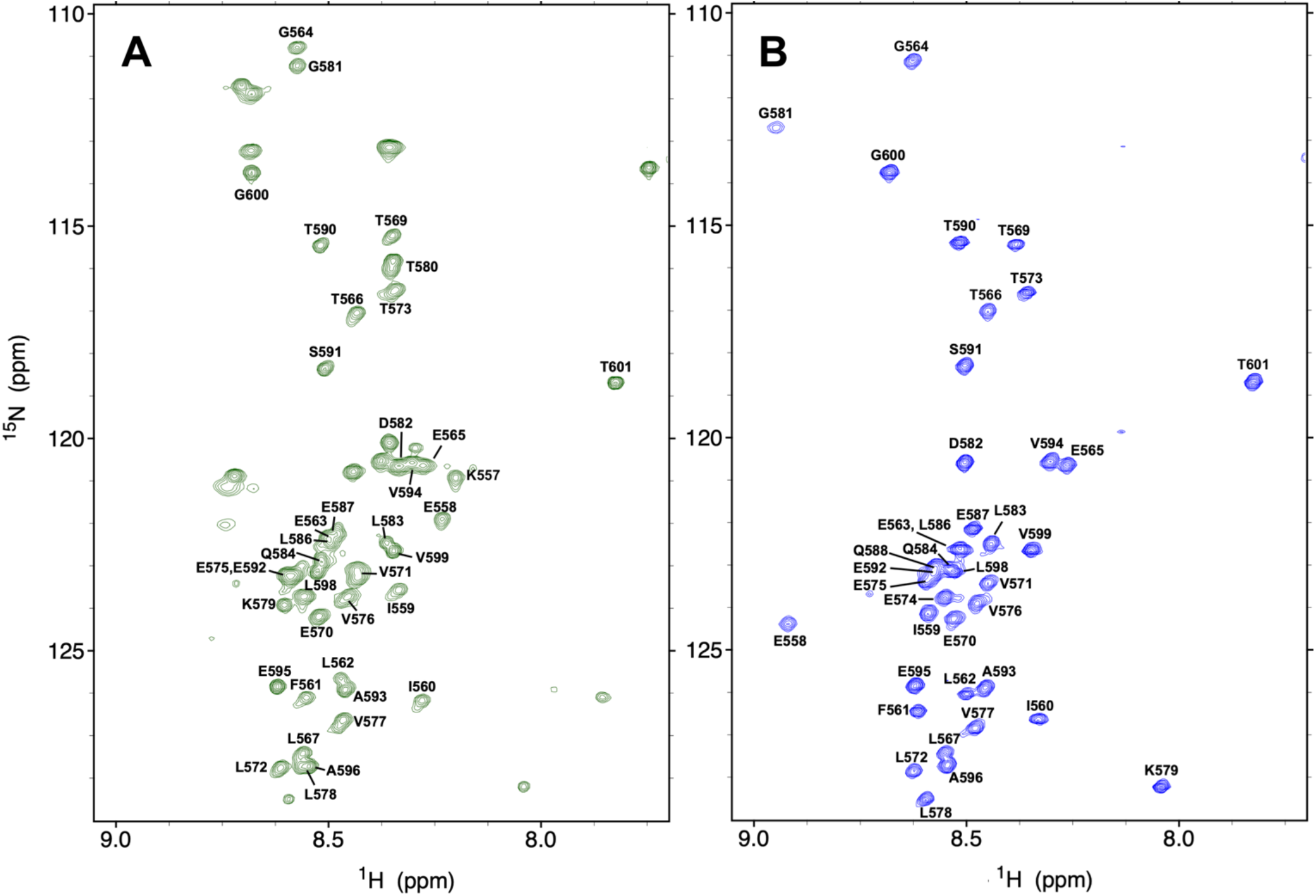
2D ^1^H-^15^N HSQC NMR Spectra of Nsp2-CtDR in 5 mM KPi, 10 mM NaCl, pH 6.3, 5°C **A.** Prior to cleavage of the His tag. Peaks are shown in **green** and residues are labeled according to their numbering in the full length Nsp2 protein. Note that peaks arising from the N-terminal His tag are not labeled: The His tag sequence is: MAHHHHHHGTGTGSNDDDD-K **B.** Following His tag cleavage; signals are shown in **blue**. Here the polypeptide contains an additional break between K_579_ and T_580_.

**Figure 3.**
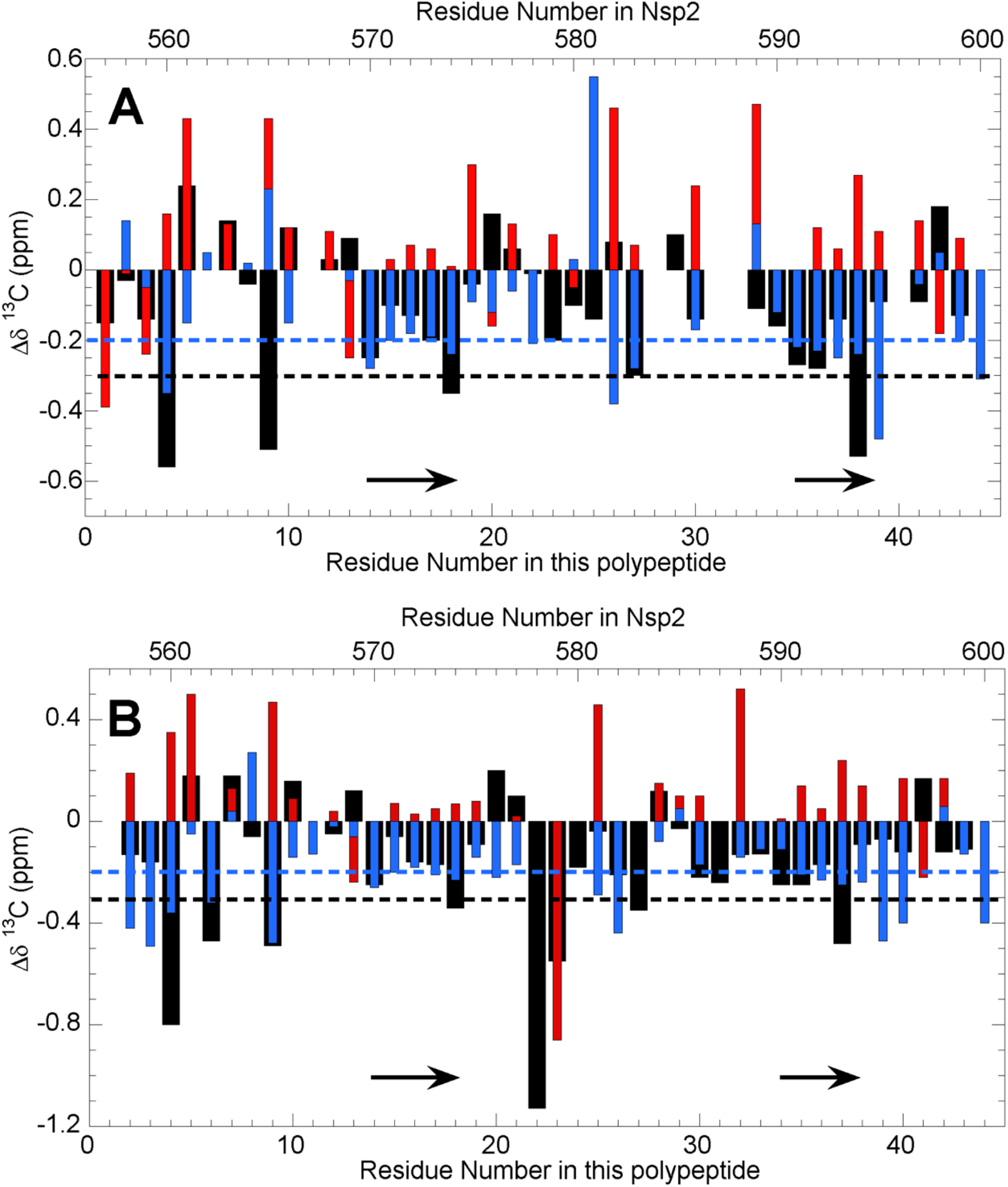
Conformational shift analysis for Nsp2-CtDR in 5 mM KPi, 10 mM NaCl, pH 6.3, 5°C Nsp2 CtDR conformational chemical shifts of ^13^Cα (**black)**, ^13^CO **(blue)**and ^13^Cβ (**red**): **A**. prior to cleaving the His tag, and **B**. after removing the His tag. Note the large chemical shift perturbations around residue T_580_ which reflects the cleavage at the K_579_-T_580_ peptide bond. The dashed **blue** and **black** lines mark the values expected for 10% β-strand for ^13^CO and ^13^Cα, respectively. The position of two modestly populated β-strands are marked by arrows.

### 3. Dynamics

Disordered proteins are often somewhat rigid; this helps predispose them for interactions with other molecules and to adopt specific conformations. To assess the μs-ms dynamics, we measured transverse relaxation rates in the rotating time frame (R_1ρ_) for the Nsp2 CtDR. Generally low values are found which reflect flexibility (**Figure 4A**). The presence of the His tag and the lack of cleavage at the K_579_-T_580_ site appear to stiffen the polypeptide somewhat. Higher than average values are detected for residues E_570_-E_574_ and S_591_-E_595_ which adopt a small β-strand population.

**Figure 4:**
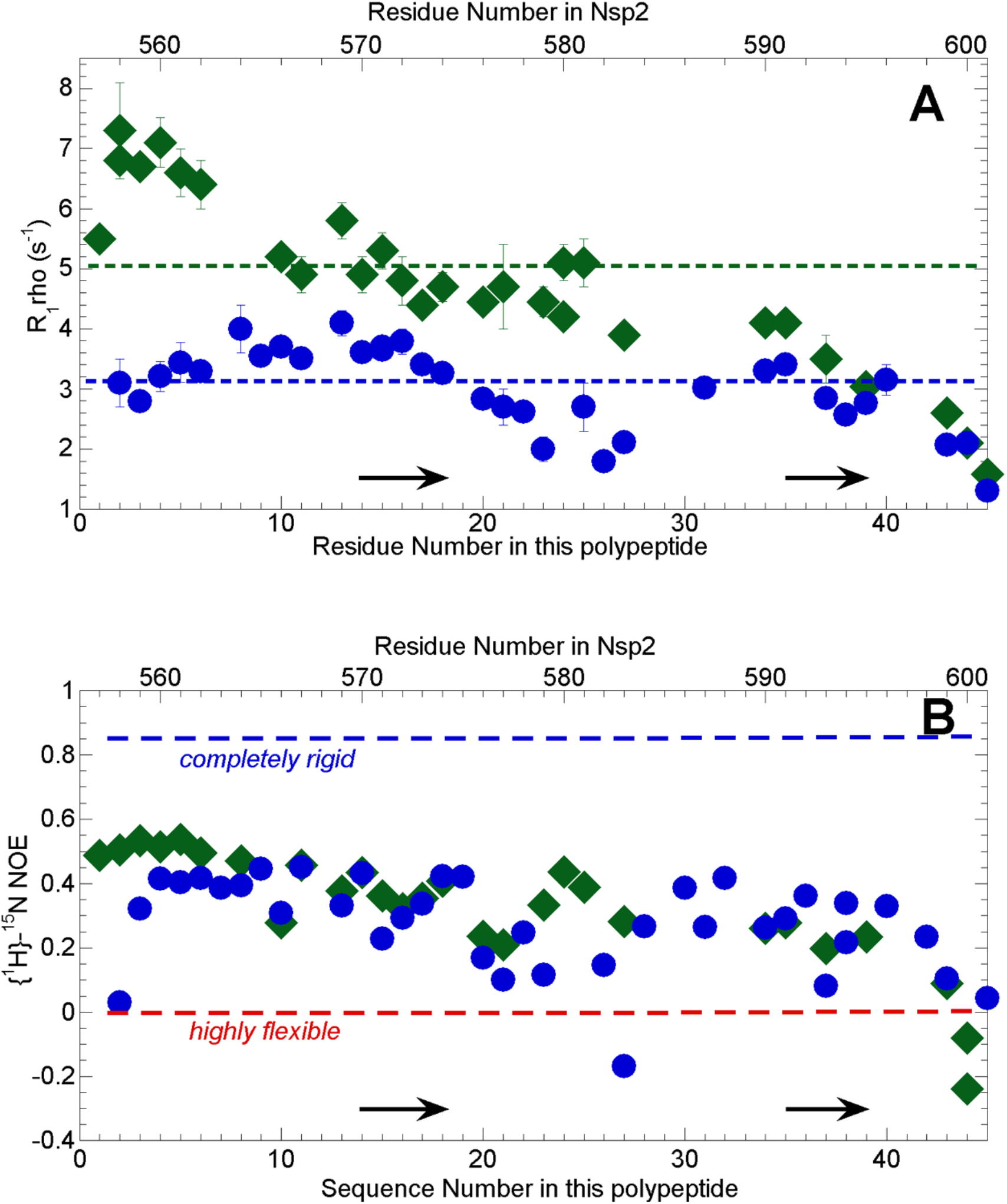
Residue Level Dynamics from ^15^N Relaxation. **A.** Transverse relaxation rates in the rotating frame (R_1ρ_) for the Nsp2 CtDR with (**green**) and without (**blue**) the His tag present. The mean values are represented by the dashed lines. The arrows show the position of the modestly populated β-strands. **B.** {^1^H}-^15^N NOE values for the Nsp2 CtDR, with (**green diamonds**) or without (**blue circles**) the N-terminal His tag. The **blue** and **red** dashed lines mark values indicative of fully rigid and highly flexible polypeptides. The arrows show the position of the modestly populated β-strands. As the calculated errores are less than 0.02, no error bars are shown as they are smaller than the symbol.

Regarding the ps-ns timescale dynamics, {^1^H}-^15^N NOE measurements show ratios about halfway between those indicative of full rigidity and complete flexibility (**Figure 4B**). Thus, the Nsp2 CtDR is significantly more rigid than fully unfolded proteins.

## Discussion

In this communication, we show that the C-terminal region of SARS-CoV-2 of Nsp2 is intrinsically disordered, which is in general agreement with bioinformatics predictions (Giri *et al.*, 2020). Nevertheless, two five-residue stretches show a small, but significant tendency to adopt β-strand conformations, and the region is more rigid than a completely disordered protein chain. These findings may be attributed to the content of aliphatic residues: two Ile, four Pro, five Val and seven Leu, which is rather high for a disordered region. These results seem not to depend on the presence of an N-terminal His tag or an internal proteolytic cleavage. Nevertheless, additional experiments on the Nsp2 intact CtDR without the His tag, which are underway, are necessary to confirm these findings. These β-strands may participate in interactions with human proteins and future experiments will test whether their degree of structure and rigidity increase upon binding. Blocking such an interaction might be an excellent target for future farmaceutics. Inhibitor screening of the Nsp2 CtDR in collaboration with the COVID19-NMR consortium is starting. In the future, the assigned ^1^H-^15^N spectrum could serve to identify where first-generation inhibitors bind and thereby help guide their improvement.

## Acknowledgements

M. Mompeán is a “Caixa” Foundation Junior Group Leader. Funded by project COV20/00764 from the Carlos III Institute of Health and the Spanish Ministry of Science and Innovation. NMR experiments were performed in the “Manuel Rico” NMR Laboratory (LMR) of the Spanish National Research Council (CSIC), a node of the Spanish Large-Scale National Facility for Biomolecular NMR (ICTS R-LRB). Inhibitor screening is being conducted within the COVID19-NMR consortium by Dr. S. Sreeramula, Prof. Dr. H. Schwalbe and co-workers.

